# Single-Droplet Surface-Enhanced Raman Scattering Decodes the Molecular Language of Liquid-Liquid Phase Separation

**DOI:** 10.1101/2022.02.05.479225

**Authors:** Anamika Avni, Ashish Joshi, Anuja Walimbe, Swastik G. Pattanashetty, Samrat Mukhopadhyay

**Author notes:** Corresponding author Samrat Mukhopadhyay, Indian Institute of Science Education and Research (IISER) Mohali, Sector 81, Knowledge, City, S.A.S. Nagar, Mohali 140306, Punjab, India. Contributed equally (Anamika Avni, Ashish Joshi).

## Abstract

Biomolecular condensates formed via liquid-liquid phase separation (LLPS) are involved in a myriad of critical cellular functions and debilitating neurodegenerative diseases. Elucidating the role of intrinsic disorder and conformational heterogeneity of intrinsically disordered proteins/regions (IDPs/IDRs) in these phase-separated membrane-less organelles is crucial to understanding the mechanism of formation and regulation of biomolecular condensates. Here we introduce a unique single-droplet surface-enhanced Raman scattering (SERS) methodology that utilizes surface-engineered, plasmonic, metal nanoparticles to unveil the inner workings of mesoscopic liquid droplets of Fused in Sarcoma (FUS) in the absence and presence of RNA. These highly sensitive measurements offer unprecedented sensitivity to capture the crucial interactions, conformational heterogeneity, and structural distributions within the condensed phase in a droplet-by-droplet manner. Such an ultra-sensitive single-droplet vibrational methodology can serve as a potent tool to decipher the key molecular drivers of biological phase transitions of a wide range of biomolecular condensates involved in physiology and disease.

## Introduction

Biomolecular condensation via liquid-liquid phase separation (LLPS) offers an exquisite mechanism for spatiotemporally-controlled organization and compartmentalization of cellular constituents into highly dynamic, permeable, liquid-like, tunable, mesoscopic, non-stoichiometric supramolecular assemblies known as membrane-less organelles^1-10^. These on-demand non-canonical organelles containing proteins and nucleic acids are present both in the cytoplasm and nucleus and include nucleoli, stress granules, P granules, nuclear speckles, and so forth. A growing body of exciting research suggests that highly flexible intrinsically disordered proteins/regions (IDPs/IDRs) containing low-complexity regions and prion-like domains that offer conformational heterogeneity, distributions, and multivalency are excellent candidates for intracellular phase separation. A unique combination of these sequence-dependent features governs the making and breaking of promiscuous and ephemeral intermolecular interactions such as electrostatic, hydrophobic, hydrogen bonding, dipole-dipole, π-π, and cation-π interactions resulting in liquid-like behavior^3,6,11-17^. While these highly dynamic condensates are proposed to be involved in a range of critical cellular functions, their transitions into less dynamic gel-like or solid-like aggregates containing more persistent interchain interactions are linked to debilitating neurodegenerative diseases. Therefore, it is imperative to decipher the molecular language of phase transitions involved in functions and disease^18^. A multitude of spectroscopic and microscopic methodologies have been employed to unveil the key biophysical principles of phase separation resulting in the formation of liquid droplets. For instance, high-resolution microscopic tools such as confocal, super-resolution, and high-speed atomic force microscopy can directly probe the properties within individual liquid droplets^19,20^. However, these tools do not allow us to access the wealth of molecular information in a residue-specific manner. On the contrary, the high-resolution structural methods such as nuclear magnetic resonance (NMR) and small-angle X-ray scattering (SAXS) provide the atomic-resolution details of the condensed phase architecture^21-23^. However, these ensemble structural methods are not capable of yielding molecular insights from the condensed phase of individual droplets. Therefore, a method that combines these capabilities to capture residue-specific structural information at a single-droplet resolution is essential to characterize and quantify the key molecular determinants in a droplet-by-droplet manner.

Vibrational Raman spectroscopy performed in a microscopy format allows us to uniquely and elegantly combine the aforesaid capabilities to obtain the protein structural information from a well-defined spatial location by focusing the laser beam into a sub-micron spot. Such non-invasive and label-free laser micro-Raman measurements permit us to access the wealth of structural information by monitoring a range of bond vibrational frequencies while retaining the spatial resolution^24,25,26^. However, owing to a low Raman scattering cross-section, Raman spectroscopy is a highly insensitive technique, especially for biomolecules under physiological conditions in aqueous solutions^27^. Additionally, a high laser power required for Raman spectroscopic detection can be detrimental for soft biological samples. The low-sensitivity issue in Raman scattering can be circumvented by a near-field plasmonic enhancement by metallic nanostructured substrates giving rise to high electromagnetic/chemical enhancement of Raman signals even at extremely low analyte concentrations. This surface-sensitive technique is known as surface-enhanced Raman scattering (SERS) that can provide several orders of magnitude increase in the Raman scattering cross-section allowing single-molecule detection and characterization even at a much lower laser power^28-32^. In the present work, we have developed an ultra-sensitive single-droplet SERS methodology that can illuminate the unique molecular details of the polypeptide chains within individual phase-separated protein liquid droplets. For our studies, we have used Fused in Sarcoma (FUS) that is one of the most intensely studied RNA-binding proteins containing archetypal prion-like low-complexity domains and hence one of the best prototypes of phase-separating proteins. The human genome encodes approximately 30 FUS-family proteins that are known to be involved in critical functions such as mRNA splicing, DNA damage repair, formation of stress granules as well as in deadly neurodegenerative diseases such as ALS (amyotrophic lateral sclerosis) and FTD (frontotemporal dementia)^33-38^. Here we show that upon liquid phase condensation, surface-coated, SERS-active, nanosphere substrates get spontaneously encapsulated within the protein-rich condensed phase and generate plasmonic hotspots that permit us to capture the inscrutable workings of FUS condensates with unprecedented sensitivity in the absence and the presence of RNA.

## Results

### Experimental design for single-droplet vibrational Raman spectroscopy

Our laser micro-Raman system consists of several components namely, an excitation source comprising of a near-infrared (NIR) laser, an integrated microscope spectrometer consisting of a combined system of lenses, mirrors, filters, and a diffraction grating, and a charge-coupled device (CCD) detector (Fig. 1). This integrated optical setup allows us to irradiate the sample and filter out the (elastic) Rayleigh scattered light and further collimate the (inelastic) Raman scattered light onto the detector to obtain a Raman spectrum. Such a design permits us to focus the laser beam of a suitable power using an objective lens into a small sub-micron spot-size within a single protein-rich droplet and acquire (regular) normal single-droplet Raman spectra. For ultra-sensitive SERS measurements, we observed that surface-modified metal nanoparticles get spontaneously encapsulated into liquid droplets as evident by an independent confocal fluorescence imaging experiment (Fig. 1). The single-droplet SERS methodology allows us to obtain highly enhanced Raman signals within individual liquid droplets of FUS.

**Fig. 1.**
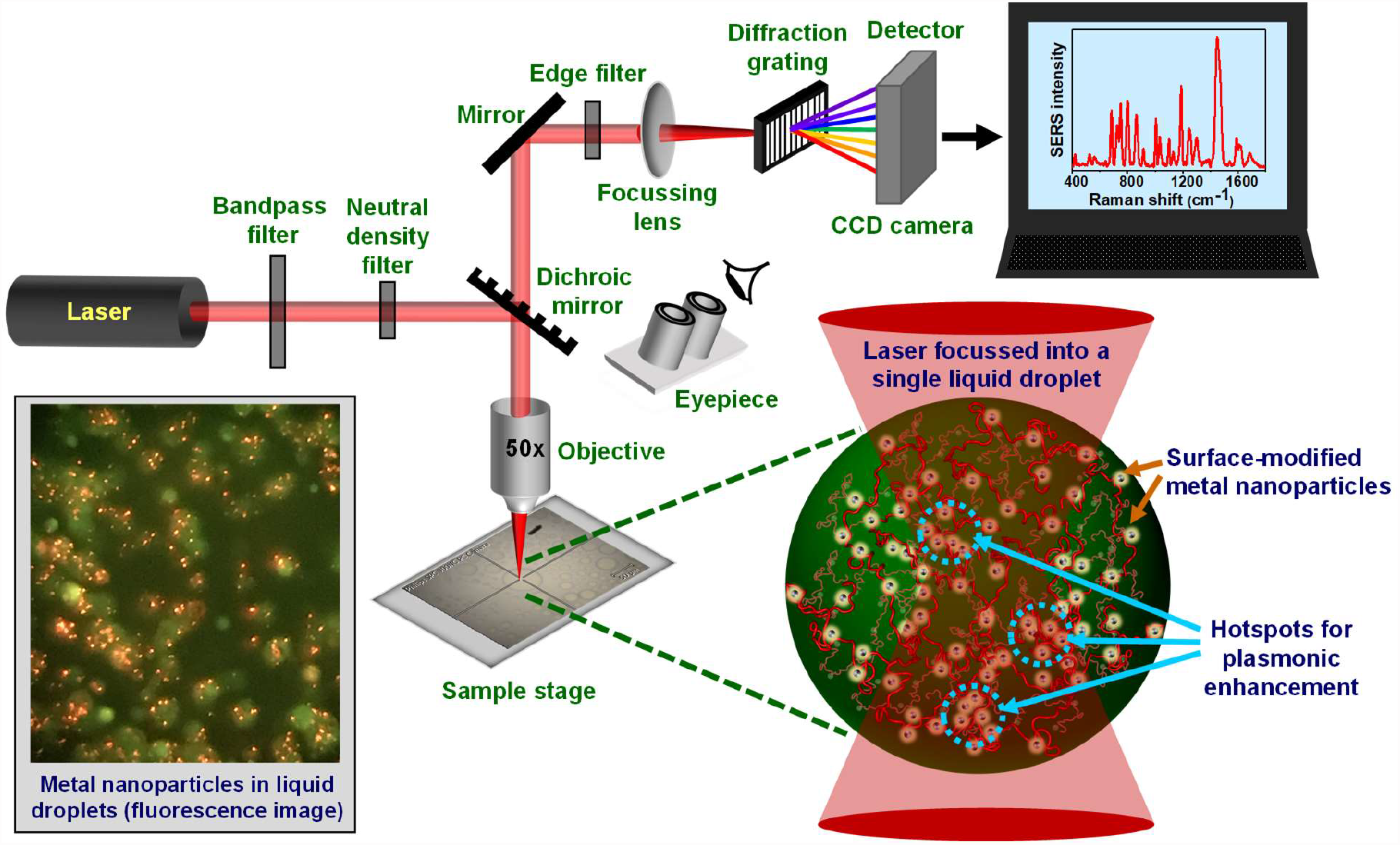
A sketch of the optical setup and a schematic of single-droplet normal and surface-enhanced Raman spectroscopy (SERS). A near-infrared (NIR) laser gets focussed within each protein-rich FUS droplet encapsulating surface-functionalized silver nanoparticles through an integrated system of lenses, mirrors, and filters. Hotspots are generated within the droplets causing optical enhancement of Raman signals detected by a CCD detector. A fluorescence image taken from the eyepiece using a camera is also included to show the encapsulation of nanoparticles in the condensates.

### Single-droplet normal Raman spectroscopy of FUS condensates

FUS consists of an intrinsically disordered N-terminal low-complexity (LC) prion-like domain (residues 1-163) and a C-terminal RNA-binding domain (RBD) containing both disordered and *α*-helical secondary structural elements (residues 267-526) (Fig. 2). The RBD contains two RGG-rich stretches, an RNA-recognition motif (RRM), and a zinc finger domain and carries a net-positive charge (+10.8) at physiological pH. The intrinsically disordered LC-domain is the primary driver of phase separation, and the presence of the RBD greatly enhances the propensity for phase transitions under physiological conditions^14,34-36^. We recombinantly expressed FUS with a cleavable N-terminal maltose-binding protein (MBP) tag as described earlier^37^. We then induced LLPS by cleaving the MBP tag using TEV protease that resulted in the condensation of a homogeneously mixed aqueous solution to liquid droplets as reported previously^37^. The condensed droplet phase was devoid of cleaved MBP as observed before (Supplementary Fig. 1)^37^. Next, we set out to perform single-droplet vibrational Raman experiments on these FUS liquid droplets by focusing the laser beam within individual liquid droplets, one at a time.

**Fig. 2.**
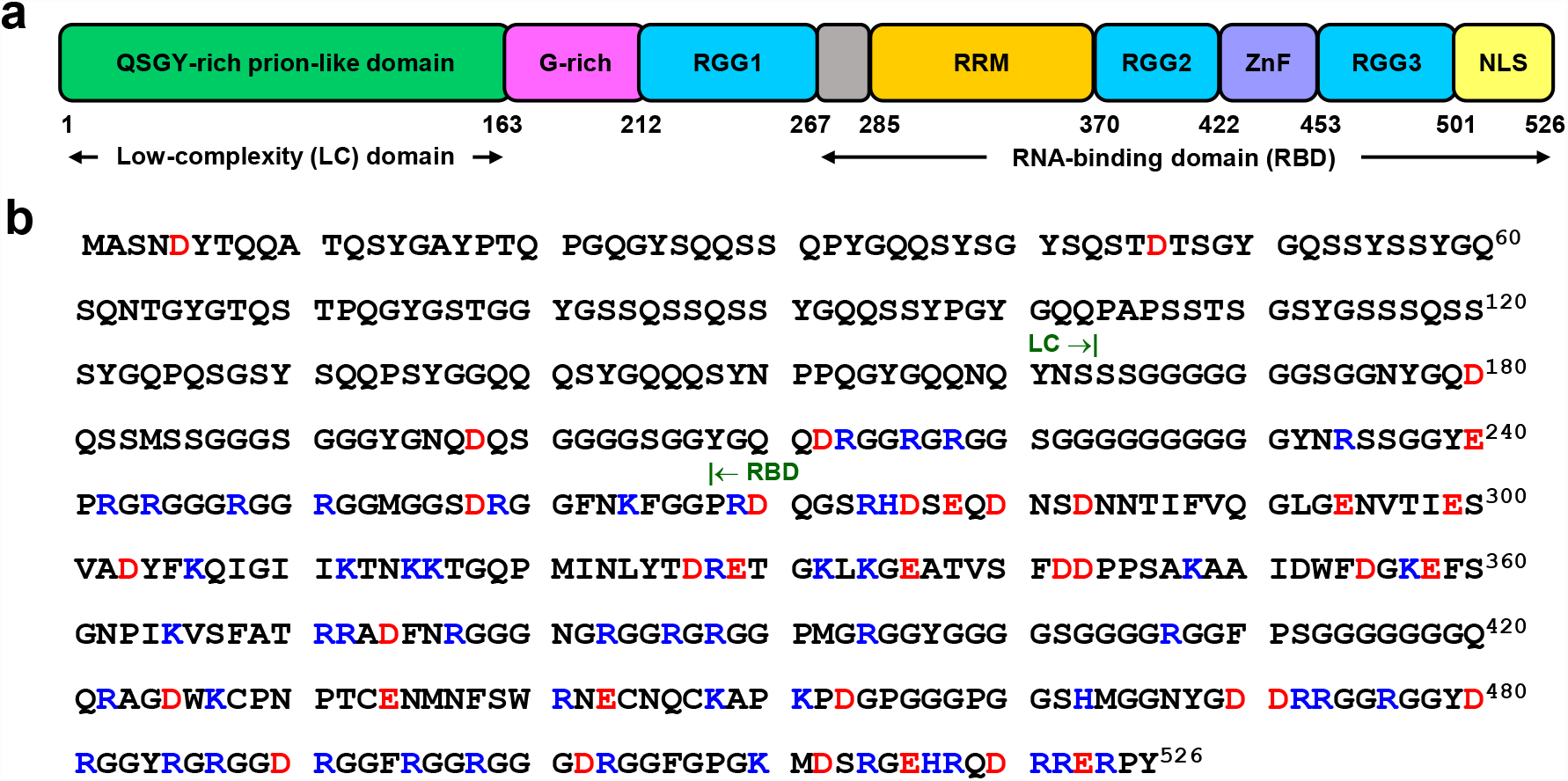
Domain architecture and amino acid sequence of full-length FUS. a, Schematic representation of full-length FUS showing all the segments and domains. b, Amino acid sequence of full-length FUS. The positively and negatively charged residues are highlighted in blue and red, respectively.

Prior to performing more advanced and involved SERS experiments, we carried out (regular) normal single-droplet Raman spectroscopy by focusing a high-power (500 mW) 785 nm laser beam using a 100x objective. These studies allowed us to characterize the condensed phase by recording the Raman scattering bands for different vibrational modes of the polypeptide chains in a droplet-by-droplet manner (Fig. 3a). Single-droplet Raman spectra were dominated by amide I (1620-1700 cm^-1^), amide III (1230-1300 cm^-1^) as well as bands due to different vibrational modes of aromatic amino acids (phenylalanine, tyrosine, and tryptophan) in addition to several other aliphatic sidechain vibrations (Fig. 3b, Supplementary Table 2)^27,39-41^. Amide I (1620-1700 cm^-1^) arise primarily due to C=O stretching vibrations and amide III (1230-1300 cm^-1^) represents C-N stretching and N-H bending vibrations of the polypeptide backbone. These amide bands in a protein Raman spectrum are the typical marker bands for secondary structural elements^42^. The condensed phase showed a broad amide I band centered at ∼ 1671 cm^-1^ with a full-width at half maximum (FWHM) ∼ 59 cm^-1^ representing a considerable conformational distribution. This was further supported by the amide III band at ~ 1262 cm^-1^ representing highly disordered conformers within the condensed phase. Further, to decode the sidechain environment, we inspected the intensity ratio at 850 cm^-1^ and 830 cm^-1^ (I_850_/I_830_) of the tyrosine Fermi doublet that is observed due to Fermi resonance between the ring-breathing vibration and overtone of an out-of-plane ring-bending vibration of the phenolic ring of tyrosine. Therefore, this ratio is an indicator of solvent-mediated hydrogen bonding propensity of the phenolic (-OH) group and is a measure of the water accessibility of tyrosine residues^43^. The I_850_/I_830_ ratio is typically ≥ 2 for a well-solvated tyrosine and this ratio was found to be ∼ 0.5 for FUS droplets indicating considerable solvent protection possibly due to the participation of tyrosine residues in π-π stacking and/or cation-π interactions in the dense phase^44^. Another important sidechain band is the tryptophan Raman band typically observed at ~ 880 cm^-1^ that arises due to the indole N-H bending and is often used to probe the environment and is a measure of the hydrogen bonding strength between the N-H of the indole ring with the surrounding solvent molecules. This band is highly blue-shifted to 891 cm^-1^ in droplets indicating a reduced hydrogen bonding propensity of the N-H group with the surrounding water molecules implying an apolar microenvironment in the vicinity of tryptophan residues^45^. Additionally, we observed a tryptophan band at 767 cm^-1^ that corresponds to the indole ring breathing and is used as a marker for cation-π/CH-π interactions^46,47^. Therefore, these observations might potentially indicate the presence of π-π/cation-π interactions within FUS condensates. We also observed a broad band at around 540 cm^-1^ corresponding to characteristic backbone deformations due to the presence of a large number of highly flexible glycine residues in FUS^48,49^. Taken together, this set of normal single-droplet Raman experiments indicate conformational heterogeneity and intrinsic disorder of FUS within the droplets. These results also revealed the involvement of aromatic sidechains of tyrosine and tryptophan residues in the chain collapse and condensation of FUS corroborating previous findings^23^. Next, in order to enhance the sensitivity as well as to detect and characterize weaker and hidden vibrational signatures in FUS and FUS-RNA condensates, we set out to perform single-droplet SERS measurements.

**Fig. 3.**
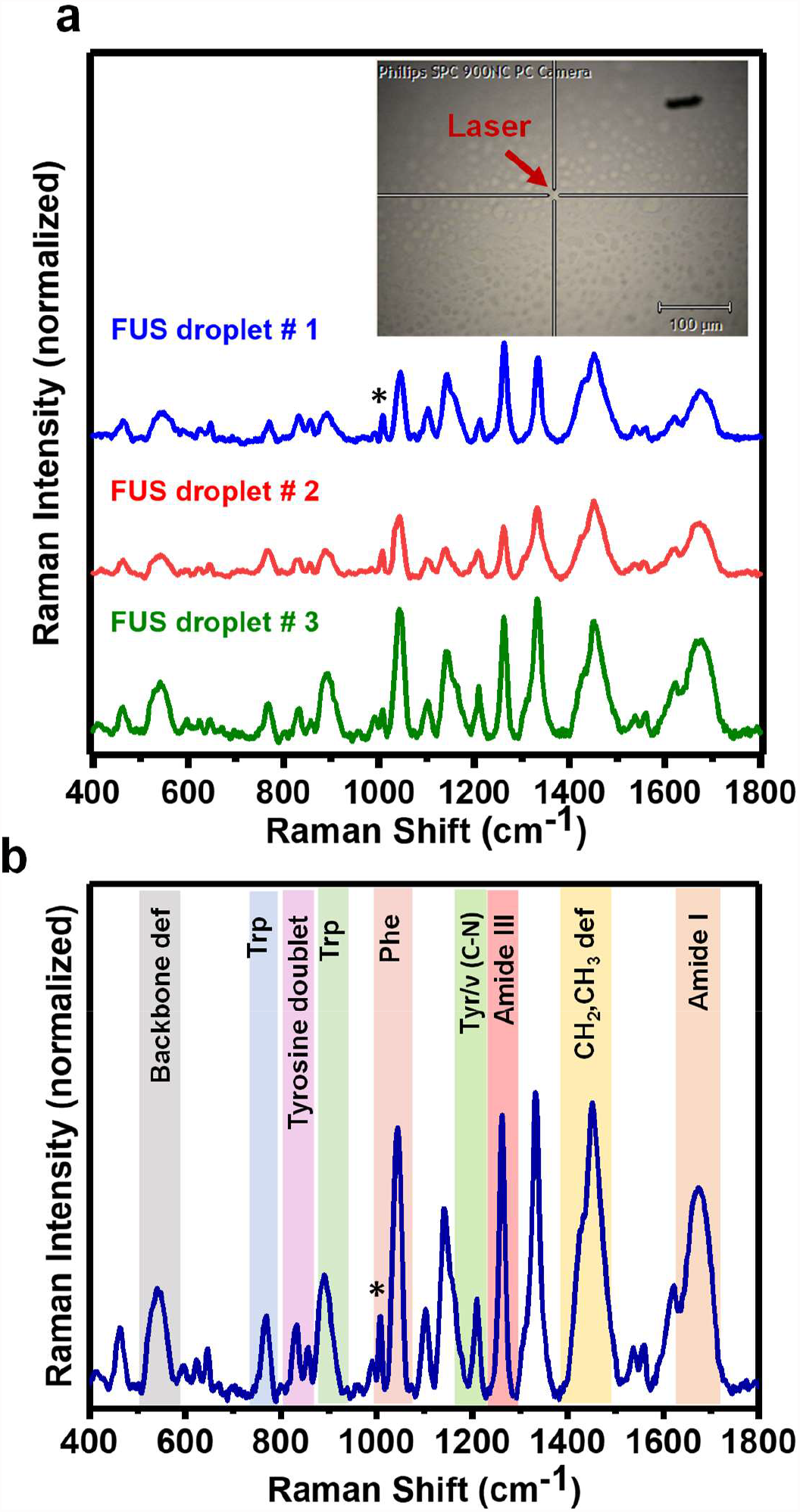
Single-droplet normal Raman spectroscopy. a, Representative single-droplet normal Raman spectra of a few individual FUS droplets (spectra recorded at 500 mW laser power, 100x objective; number of droplets, n = 3). Inset shows single-droplets of FUS focussed through the Raman microscope. Arrowhead shows the focal point of the NIR laser within the droplet. b, Average Raman spectrum of the FUS condensed phase. All spectra are normalized with respect to the phenylalanine ring breathing band at 1007 cm^-1^ marked by an asterisk. See Methods for details of data acquisition, processing, and analysis.

### Preparation of surface-modified nanoparticles for single-droplet SERS

The predicted net charge of FUS is +14.5 at physiological pH and we thus postulated that it could electrostatically interact with surface-modified negatively charged metal nanoparticles offering us an excellent system to study SERS within biomolecular condensates. To test our hypothesis, we started with the preparation of a suitable plasmonic SERS substrate. We chose silver nanoparticles (Ag NPs) for our experiments due to their high SERS activity and greater electromagnetic enhancements compared to other plasmonic nanomaterials. We prepared Ag NPs by a standard protocol of reduction of AgNO_3_ by sodium citrate and characterized them using UV-visible absorption spectroscopy, transmission electron microscopy (TEM), and zeta (*ζ*) potential measurements (Supplementary Fig. 2)^50^. The absorption spectrum showed a single absorption band at ∼ 415 nm which corresponds to spherical nanoparticles with a diameter of ~ 30 nm (Supplementary Fig. 2a)^51^. We next functionalized these Ag NPs with iodide to form iodide-modified silver nanoparticles (Ag IMNPs). This halide modification of nanoparticles was performed to get rid of the overwhelming citrate peaks in the Raman spectrum and aid in better attachment of the polypeptide chains to the negatively charged silver nanospheres (Supplementary Fig. 2b)^50^. Ag IMNPs exhibited a single absorption band with λ _max_ at ∼ 418 nm indicating a similar diameter of surface-modified nanoparticles (∼ 30-40 nm). The size was then confirmed by TEM that revealed nanospheres with an average diameter of ∼ 30 nm (Supplementary Fig. 2c). We next carried out zeta potential measurements to determine the effective charge on the surface of nanoparticles. The zeta potential of Ag IMNPs was −22 mV indicating an overall negative surface charge that stabilizes the nanoparticles preventing them from agglomerating into large-sized colloids (Supplementary Fig. 2d). Further, Raman spectra of these surface-modified nanoparticles showed a single band at 110 cm^-1^ that corresponds to Ag-I bond indicating monolayer coating of Ag NPs (Supplementary Fig. 2e)^50^.

We next set out to investigate the interaction of FUS with Ag IMNPs and to check their colloidal stability in the presence of the protein. We recorded UV-vis absorption spectra of Ag IMNPs in the dispersed and condensed phase of FUS at 10 and 30 minutes time points and observed a small red-shift of the absorption maxima with a slight broadening of the band as compared to only Ag IMNPs in buffer (Supplementary Fig. 3a,b). We chose to record the absorption spectra at these two different time points since our next set of single-droplet SERS measurements required this time interval for data acquisition. Absorption data indicated that positively charged FUS electrostatically binds to the negatively charged nanoparticle surface in the monomeric and the phase-separated state without altering the stability of nanoparticles^52^. A slight red-shift also suggested that the presence of FUS allowed the clustering of nanoparticles to create critical hotspots necessary for plasmonic enhancements in Raman measurements. In order to test if the LLPS behavior of FUS remained unaltered in the presence of nanoparticles, we performed turbidity and microscopy assays that indicated nanoparticles do not alter the phase separation propensity of FUS (Supplementary Fig. 3c,d,e). To visualize the presence of nanoparticles within the condensates, we performed two-color confocal fluorescence imaging that revealed the uptake of nanoparticles within the condensates (Fig. 4a,b and Movie S1). Fluorescence recovery after photobleaching (FRAP) experiments on fluorescently-labeled FUS droplets revealed no significant difference in the rate of recovery in the absence and the presence of nanoparticles (Fig. 4c). These results indicated that the droplet interior remained mobile in the presence of nanoparticles and the overall material property of FUS condensates remained unaltered in the presence of the SERS substrate. Together, this set of experiments suggested that FUS electrostatically interacts with surface-coated silver nanoparticles that get preferentially encapsulated into the dense phase of liquid droplets while keeping the internal mobility nearly intact. Therefore, these silver nanoparticles can act as an ideal SERS substrate for Raman enhancements within biomolecular condensates. We next directed our efforts to perform ultrasensitive SERS measurements within individual droplets.

**Fig. 4.**
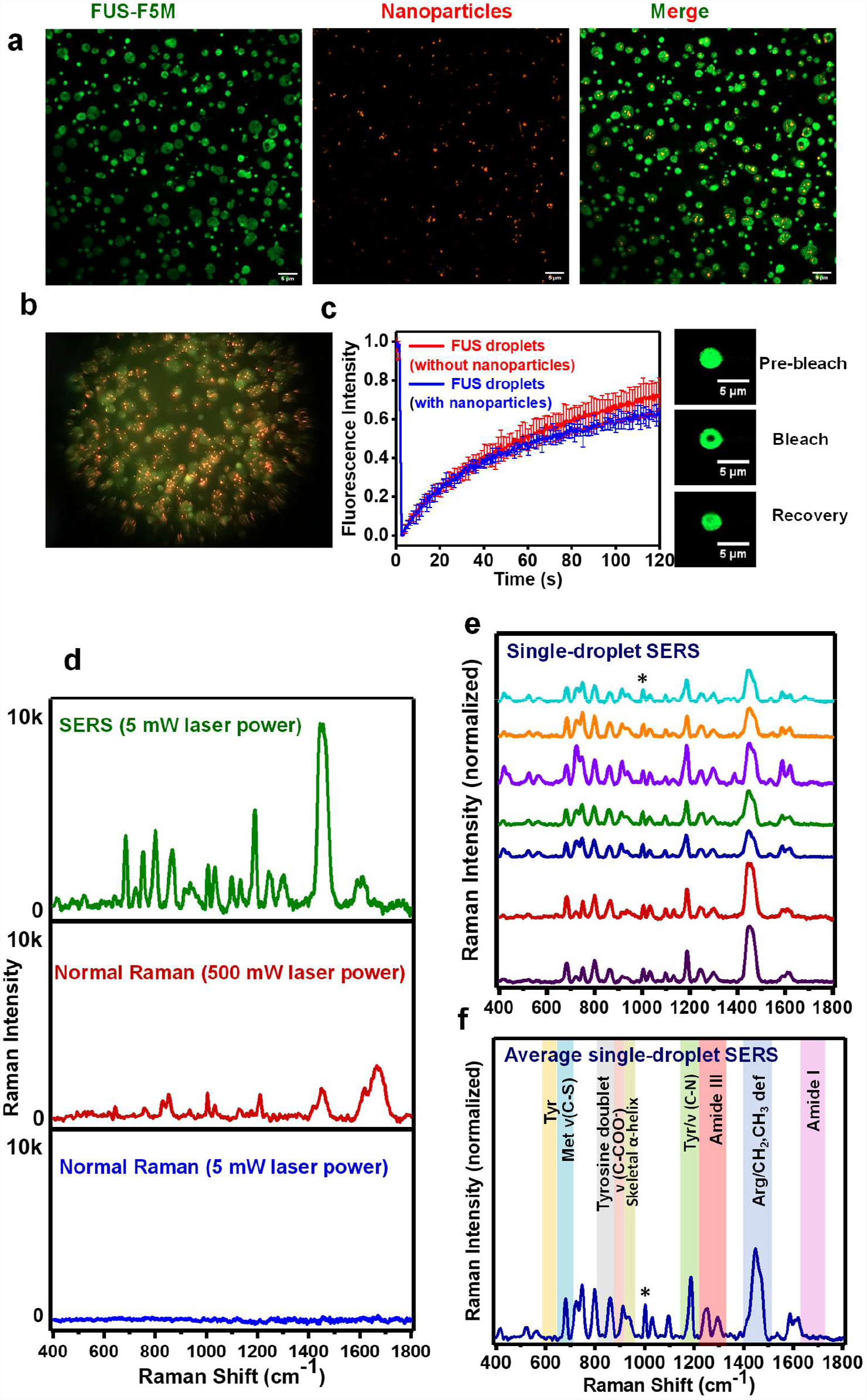
Spontaneous encapsulation of iodide-modified silver nanoparticles (Ag IMNPs) in FUS liquid droplets and single-droplet SERS. a, Confocal images depicting encapsulation of iodide-modified silver nanoparticles (Ag IMNPs) within fluorescein-5-maleimide labeled FUS droplets. See Movie 1 for 3D Z-stack images of droplets containing nanoparticles. b, Image clicked through the eyepiece using a camera indicating the encapsulation of Ag IMNPs within FUS droplets (a similar image is also shown in a scheme in Fig. 1). c, FRAP kinetics of multiple droplets (∼1% Alexa488-labeled protein; n = 3) in the absence (red) and presence of nanoparticles (blue). Fluorescence images of droplets during FRAP measurements are shown on the right. d, Single-droplet SERS (5 mW laser power), single-droplet normal Raman (500 mW laser power), and single-droplet normal Raman spectra (5 mW laser power) using 50x objective lens. The observed enhancement was of the order of ≥ 10^4^ times using Amide III as a reference peak. e, Stacked single-droplet SERS spectra of 7 FUS droplets in the presence of Ag IMNPs (spectra recorded at 5 mW laser power with a 50x objective). f, Average single-droplet SERS spectra from individual droplets encapsulating Ag IMNPs (n = 7). See Methods for experimental details and Supplementary Table 2 for all the band positions. All spectra are normalized with respect to the phenylalanine ring breathing band at 1003 cm^-1^ marked by an asterisk. See Methods for details of data acquisition, processing, and analysis.

### Single-droplet SERS within FUS condensates

In order to record single-droplet SERS spectra, LLPS was set up in the presence of 100 pM Ag IMNPs, and a 785-nm NIR laser beam (5 mW) was focused into individual nanoparticle-containing liquid droplets using a 50x objective (Supplementary Fig. 3f). We achieved an enhancement in the order of ≥ 10^4^ using amide III as a reference peak. We would like to note that this is an approximate (lower bound) estimate of the enhancement factor due to extremely weak signals from the droplets in the absence of nanoparticles using low power and a 50x objective (Fig. 4d). The enhancement is much higher than 10^4^ for the peaks that are not visible in normal Raman. Interestingly, the amide I band in our SERS spectra was not enhanced, and therefore, was not visible in such a low power illumination (Fig. 4e,f). According to selection rules, signal enhancement in SERS depends on the orientation of the analyte on the surface of nanoparticles and varies inversely with the twelfth power of the distance of the target analyte from the nanoparticle surface^53^. We speculate that the interaction of bulky amino acid side chains of FUS with the negatively charged Ag IMNPs might potentially orient the backbone C=O group away from the critical near-field required for the plasmonic enhancement^54^. Nevertheless, amide III was visible at 1246 cm^-1^ and 1298 cm^-1^ corresponding to nonregular/turn structures and α-helices, respectively (Fig. 4e,f). Several orders of magnitude signal enhancement allowed us to observe these structures and is possibly caused by electrostatic interactions between the negatively charged surface of Ag IMNPs and arginine-rich positively charged RBD containing these secondary structural elements. In addition, a significant enhancement was also observed for backbone C-C stretch of α-helices at 938 cm^-1^. Surprisingly, closer inspection of the tyrosine Fermi doublet showed that the lower wavenumber band at 830 cm^-1^ did not show significant enhancement while the higher wavenumber band showed enhancement and shifted to 863 cm^-1^. This may be attributed to the SERS selection rules, according to which the polarizability component of the 830 cm^-1^ mode may not be perpendicular to the metal surface. Further, single-droplet SERS spectra were dominated by bands at 683 cm^-1^ [methionine/δ(CH)]; 724 cm^-1^ (methionine); 749 cm^-1^ (tryptophan); 915 cm^-1^ ν(COO^-^); 1003, 1032 cm^-1^ (phenylalanine); 1246, 1298 cm^-1^ (Amide III); 1588 cm^-1^ (phenylalanine/tryptophan/histidine); 1621 cm^-1^ (tyrosine) (Supplementary Table 2). Additionally, we observed a highly enhanced band at 1447 cm^-1^ and a shoulder at 1465 cm^-1^ that is assigned to the bending δ(NH) of the guanidinium moiety of arginine residues and CH_2_/CH_3_ deformation modes, respectively^55,56^. This is in accordance with the fact that the structured C-terminal RBD of FUS contains 37 arginine residues that can facilitate its adsorption to the negatively charged SERS substrate, thereby resulting in a significant enhancement in our SERS spectra. Taken together, this set of single-droplet SERS illuminated the inner conformational details within FUS condensates. We next asked whether this ultrasensitive tool can be utilized to elucidate the structural details of FUS-RNA heterotypic condensates.

### Illuminating FUS-RNA heterotypic condensates using single-droplet SERS

RNA is known to modulate the phase behavior and biophysical properties of liquid condensates formed by several RNA-binding proteins including FUS^4,57,58^. With the objective to elucidate the effect of polyU RNA on the chain conformations within the droplets, we performed normal Raman and SERS at different stoichiometries of RNA and protein (Fig. 5a,b and Supplementary Fig. 4). A careful inspection of normal single-droplet Raman spectra showed two RNA marker bands, a shoulder band at 782 cm^-1^ and a band at 1230 cm^-1^ corresponding to uracil breathing and ring stretching modes, respectively (Fig. 5c,d)^59^. This is corroborated by the Raman difference spectrum as well (Fig. 5e). An increase in the RNA concentration leads to its greater recruitment within the phase-separated droplets which is confirmed by the linear plot of peak intensity at 1230 cm^-1^ as a function of RNA concentration (Fig. 5f). Additionally, these single-droplet Raman measurements also allowed to obtain the stoichiometry of RNA and protein within condensates by following the ratio of intensities at 1230 cm^-1^ (uracil ring stretching of RNA) to at 1450 cm^-1^ (CH_2_/CH_3_ deformation modes of protein). A linear relationship obtained can be used as a calibration line to evaluate the stoichiometry of RNA and protein within the condensed phase (Fig. 5g). Such quantitative and ratiometric estimates can be valuable in determining the concentration and composition of complex multi-component and multi-phasic condensates. Upon a closer inspection of the amide I region, we observed a considerable blue shift (Fig. 5h) from 1671 cm^-1^ to 1682 cm^-1^ indicating a β→disorder conversion with an increase in RNA concentration. This could potentially be due to the formation of more liquid-like condensates having more disorder and less β-content at higher RNA concentrations. Since the uracil carbonyl (C(4)=O) stretching mode can appear around this region,^60^ we next zoomed into the amide III region to independently confirm this unraveling of FUS in the presence of RNA resulting in a more liquid-like behavior of FUS-RNA heterotypic condensates. (Fig. 5i) In this amide III region as well, in the absence of RNA, we observed primarily random-coils (∼ 1262 cm^-1^) with a small contribution of β-structure (broad shoulder at ∼ 1248 cm^-1^) that disappeared at higher RNA to protein stoichiometry. This observation also supports that the observed blue-shift in the amide I might be due to an increase in the disordered content rather than the C(4)=O stretching mode of the uracil ring. Additionally, we observed lower α-helical contents within the droplets (amide III: 1325 cm^-1^; amide I: 1660 cm^-1^) in the presence of RNA as shown in the Raman difference spectrum (Fig. 5e). This observation also hinted at a possible unwinding of the helical region of RBD. We next set out to characterize these conformational changes within the liquid condensates as a function of RNA concentration using ultra-sensitive SERS.

**Fig. 5.**
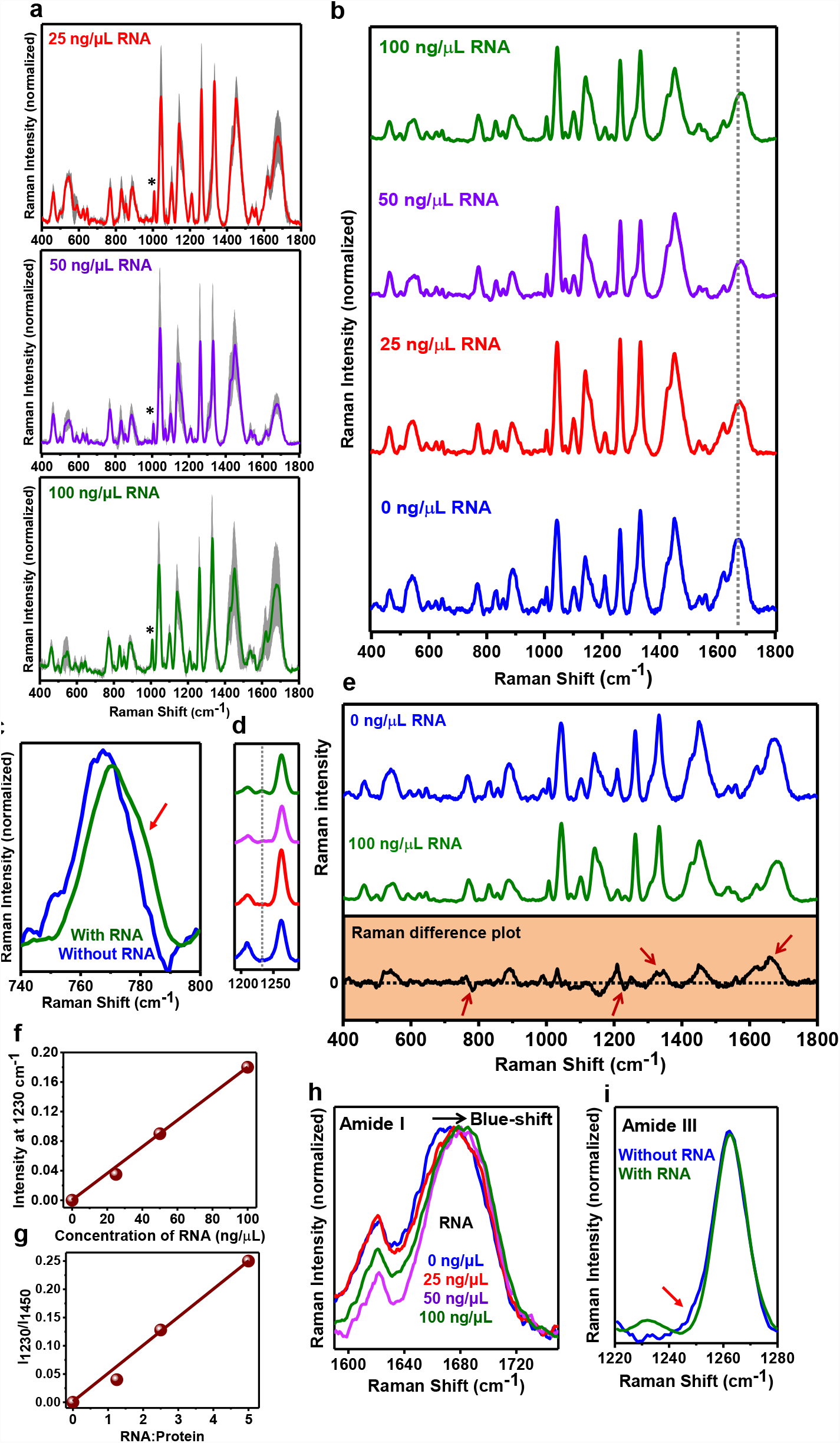
Single-droplet normal Raman spectra of FUS in the presence of RNA. a, Average single-droplet normal Raman spectra in the presence of 25 ng/μL, 50 ng/μL, and 100 ng/μL poly-U RNA (spectra recorded at 500 mW laser power with a 100x objective; number of droplets, n = 3). Solid lines represent the mean, whereas shaded region represents the standard deviation (n = 3). All spectra are normalized with respect to the phenylalanine ring breathing band at 1003 cm^-1^ marked by an asterisk. b, Stacked average single-droplet normal Raman spectra for different concentrations of RNA are shown in blue (0 ng/μL RNA), red (25 ng/μL RNA), purple (50 ng/μL RNA), and olive (100 ng/μL RNA) for comparison (the dotted line shows the shift in amide I in the presence of RNA). c, Shoulder band at ∼ 782 cm^-1^ indicated by a red arrow corresponding to the uracil ring breathing mode for FUS droplets in the presence of 100 ng/μL RNA. d, Another RNA marker band at ∼ 1230 cm^-1^ corresponding to the uracil ring stretching mode. e, Raman difference plot (between 0 ng/ μL RNA and 100 ng/ μL RNA) of single-droplet normal Raman spectra of droplets in the absence and presence of 100 ng/μL RNA (arrows indicate the differences of interest). Arrows at 780 cm^-1^ and 1230 cm^-1^ represents the RNA marker bands. Arrows at 1325 cm^-1^ and 1660 cm^-1^ denotes greater α-helical content within FUS droplets in the absence of RNA. f, Linear plot of RNA marker band at 1230 cm^-1^ versus concentration of RNA (ng/μL) used. g, Plot of ratio of intensity at 1230 cm^-1^ and 1450 cm^-1^ versus RNA: protein. h, Zoomed in amide I region for FUS droplets at different concentrations of RNA indicating blue-shift of band maxima. i, Zoomed in amide III region band for FUS droplets in the absence and presence of RNA (100 ng/μL). Arrow indicates a shoulder at ∼ 1248 cm^-1^ denoting small contribution of β-sheets in addition to random-coil structures within droplets in the absence of RNA. See Methods for details of data acquisition, processing, and analysis.

Prior to performing SERS within droplets at various concentrations of RNA, we carried UV-vis absorption spectroscopy that established the stability of nanoparticles in the presence of RNA (Supplementary Fig. 5a,b). Our turbidity and imaging assays showed that SERS substrate does not alter the behavior of FUS-RNA droplets (Supplementary Fig. 5c,d). Confocal microscopy imaging revealed complete encapsulation of nanoparticles within these droplets. We next set out to record single-droplet SERS (Fig. 6a and Supplementary Fig. 5e). Fig. 6b depicts the stacked SERS spectra from individual droplets at varying concentrations of RNA. Interestingly, we observed a broad amide I band centered at 1682 cm^-1^ that was undetected in SERS within FUS-only droplets. This amide I peak represents disordered polypeptide conformers with some of the β structures within the FUS-RNA droplets which is also depicted in the Raman difference plot (Fig. 6c). We believe that the interactions between the negatively charged phosphate backbone of RNA and positively charged C-terminal RBD alter the orientation of polypeptide chains on the surface of nanoparticles which brings the C=O groups of the polypeptide backbone in proximity to the nanoparticle surface for enhancement to occur. A closer inspection of the amide III region showed a broad band centered at 1245 cm^-1^ corresponding to β-rich and disordered/extended conformations and a band at 1300 cm^-1^ corresponding to α-helical structures for all the RNA concentrations. We observed that with an increase in RNA concentration, there is a decrease in the intensity of the amide III band at 1300 cm^-1^ implying a reduction in the overall α-helical content indicating RNA-induced structural loss in FUS-RNA condensates (Fig. 6d). Interestingly, a careful inspection of the skeletal C-C stretching mode of α-helical structures at 940 cm^-1^ showed a significant disappearance as a function of RNA (Fig. 6e). This is probably because the electrostatic interactions between the protein and RNA disrupt the proposed cation-π interactions between tyrosine residues in the LC domain and arginine residues in the RBD^14,61^. We propose that the interaction between a polyanion and the FUS increases the intrinsic disorder within the polypeptide chains at the expense of α-helical structures. Moreover, we observed changes in intensities of several vibrational modes associated with aromatic residues, tyrosine, and tryptophan at 683, 724, 749, 800, 915, 1188, 1588, 1621 cm^-1^ which indicate the changes in the orientation of the aromatic ring of these residues on the nanoparticle surface in the presence of RNA. Deconvolution and analysis of the band corresponding to N-H deformations of the guanidinium moiety of arginine and shoulder band for CH_2_/CH_3_ deformations indicated a reduction in the enhancement of arginine residues with an increase in the RNA concentration (Fig. 6f, Supplementary Table 1).

**Fig. 6.**
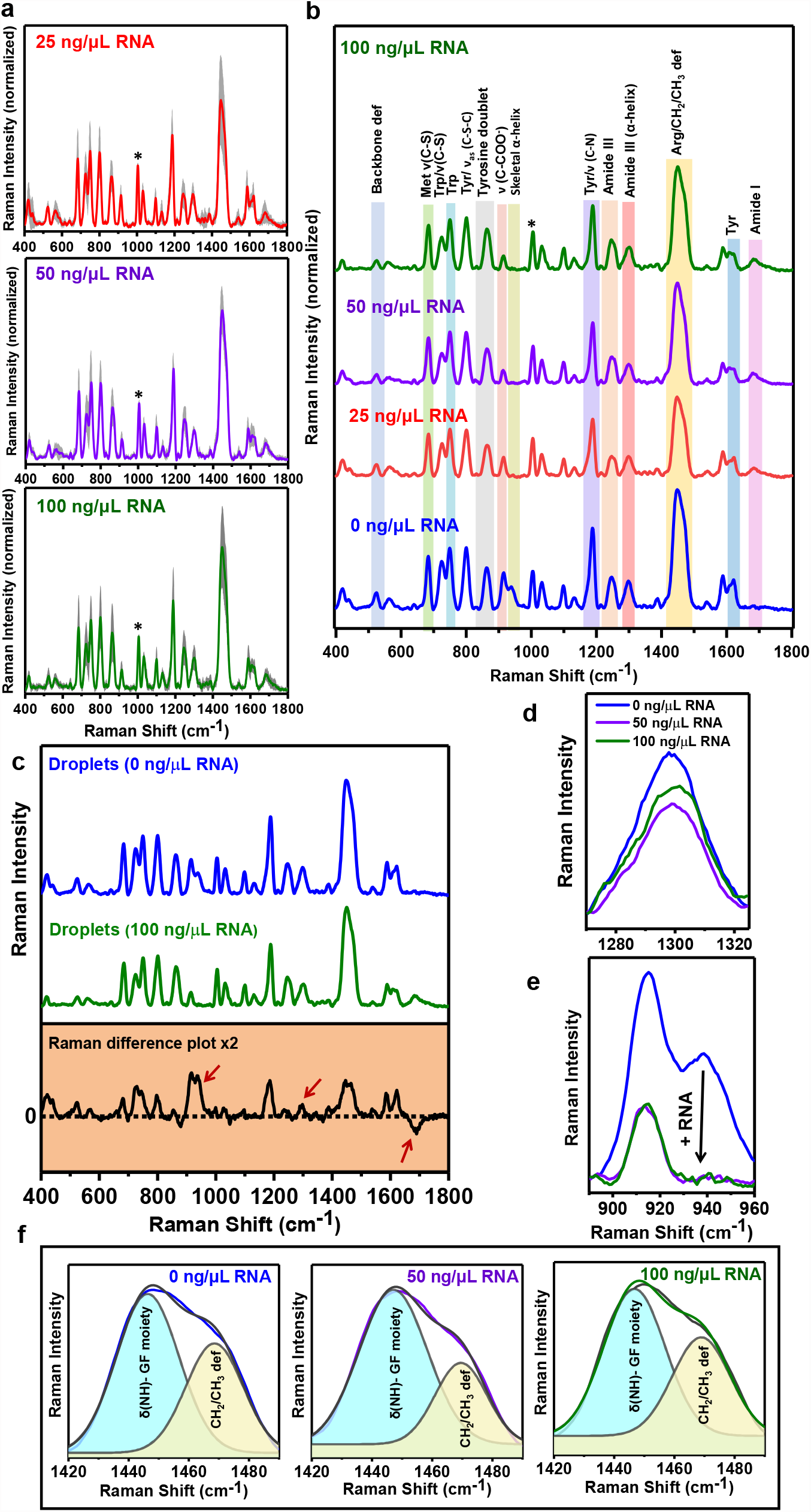
Single-droplet SERS in the presence of RNA. a, Average single-droplet SERS spectra in the presence of 25 ng/μL, 50 ng/μL, and 100 ng/μL polyU RNA (spectra recorded at 5 mW laser power with a 50x objective; number of droplets, n = 7). Solid lines represent mean, whereas shaded region represents the standard deviation (n = 7). All spectra are normalized with respect to the phenylalanine ring breathing band at 1003 cm^-1^ marked by an asterisk. b, Stacked average single-droplet SERS spectra for different concentrations of RNA are shown in blue (0 ng/μL RNA), red (25 ng/μL RNA), purple (50 ng/μL RNA), and olive (100 ng/μL RNA) for comparison. c, Raman difference plot (between 0 ng/ μL RNA and 100 ng/ μL RNA) of single-droplet SERS spectra of droplets in the absence and presence of RNA (100 ng/μL) An arrow at 1682 cm^-1^ represents the emergence of amide I at higher RNA concentrations and arrows at 940 cm^-1^ and 1300 cm^-1^ represents greater α-helical content within droplets in the absence of RNA. d, Zoomed in Amide III region for FUS droplets in the absence and presence of RNA (50 ng/μL, 100 ng/μL). e, Skeletal C-C stretching mode of α-helical structures at 940 cm^-1^ that disappears at higher RNA concentrations. f, Gaussian deconvolution of the region 1420-1490 cm^-1^. The black line represents the actual data while the colored lines represent the cumulative fit. Cyan region represents the N-H deformations of the guanidinium fragment (GF) of arginine residues, while the light-yellow region represents the CH_2_/CH_3_ deformations. See Supplementary Table 1 for percentage analysis. See Methods for details of data acquisition, processing, and analysis.

This observation directly captures the interaction between RNA and FUS by modulating the polypeptide orientation on the SERS substrate surface. Taken together, our single-droplet SERS results illuminate some key structural details within FUS-RNA condensates and highlight RNA-mediated partial unwinding of the structured domains in the C-terminal RBD.

## Discussion

We developed an ultrasensitive single-droplet Raman spectroscopic methodology to elucidate the inherent conformational heterogeneity and structural distribution within biomolecular condensates of FUS in a droplet-by-droplet manner. This unique methodology combines the capabilities of vibrational spectroscopy and optical microscopy offering a wealth of molecular information within the mesoscopic liquid condensed phase at the single-droplet resolution. Normal vibrational Raman spectroscopy can probe the detailed molecular structure and conformational reorganizations of the internal and external components of individual liquid droplets^62^. However, recording detailed vibrational signatures from liquid states is highly challenging due to a low Raman scattering cross-section of proteins^27^. Therefore, such measurements lack adequate sensitivity and often require unusually high concentrations, laser power, and magnifications. Such requirements can be detrimental to soft biological samples and lead to laser-induced damage and other artifacts. These limitations can be elegantly overcome by surface-engineered metal nanoparticle-induced plasmonic enhancements. The electrostatic interaction between positively charged polypeptide chains of FUS and negatively charged iodide-coated silver nanoparticles causes significant plasmonic enhancement of certain protein vibrational modes. By focusing a low-power laser beam into each droplet encapsulating surface-coated plasmonic nanostructures permitted us to record the Raman scattering bands arising due to different vibrational modes from the mesoscopic protein-rich droplets. We took advantage of the versatility of this technique to capture complex conformational characteristics of heterotypic FUS-RNA condensates at a single-droplet resolution.

In summary, our single-droplet Raman results showed an increase in the structural heterogeneity within liquid droplets of FUS. Several aromatic amino acid residues such as tyrosine and tryptophan residues display characteristics of the conformationally restricted environment in the condensed phase hinting at intermolecular π-π and/or cation-π interactions within liquid phase condensates. Further, this methodology allowed us to capture the unique spectral markers for droplets formed in the presence of varying RNA-protein ratios that can be used to estimate the stoichiometry of other complex biomolecular condensates of proteins and nucleic acids. The sensitivity of the single-droplet Raman methodology can be enhanced ≥ 10^4^-fold in the SERS format. Our SERS spectra showed that the C-terminal RBD undergoes a partial unwinding in the presence of RNA together with the reduction in the enhancement of arginine residues. This unraveling of the ordered region in the RBD increases the polypeptide chain disorder that can promote heterotypic interactions between FUS and RNA within the condensed phase. Therefore, taken together single-droplet SERS allows us to zoom into the mesoscopic condensed phase to unmask the molecular determinants governing the intriguing condensate biophysics. This potent methodology also offers a unique capability and adaptability by using different surface functionalities and other metals for enhancements of unique sets of vibrational bands. Additionally, cellular uptake of these engineered nanoparticles can open new avenues to study intracellular phase transitions using vibrational spectroscopy. Such advancements will pave the way for ultrasensitive detection, characterization, and quantification of a wide range of biomolecular condensates involved in physiology and disease as well as in emerging applications in drug delivery and synthetic biology.

## Methods

### Recombinant protein expression, purification

The plasmid expressing MBP-*Tev-*FUS-*Tev-*His_6_ was transformed into *E. coli* BL21(DE3) RIPL strain. The expression and purification protocols followed were similar as previously described with some slight modifications^37^. For overexpression, cultures were grown in LB media at 37 °C, 220 rpm till O.D._600_ reached 0.8-1 and was induced with 0.1 mM isopropyl-β-thiogalactopyranoside (IPTG) at 12 °C for 22 hours. Cell pellets were obtained by centrifugation at 4°C, 4000 rpm for 40 minutes, and stored at −80 °C for further use. For purification, pellets were resuspended in lysis buffer (50 mM sodium phosphate, 300 mM NaCl, 40 mM imidazole, 10 μM ZnCl_2_, 4 mM BME, and 10% v/v glycerol, pH 8.0) and cell-lysis was done using probe sonication at 5% amplitude, 15 sec ON and 10 sec OFF for 25 minutes. The lysate was centrifuged at 4 °C, 11,400 rpm for 1 hour, followed by incubation of the supernatant with equilibrated Ni-NTA agarose beads for 1.5 hours at 4 °C. The beads were washed and protein was eluted with 250 mM imidazole followed by binding to the amylose column. Protein was eluted with 20 mM maltose elution buffer (50 mM sodium phosphate, 800 mM NaCl, 40 mM imidazole, 10 μM ZnCl_2_, 20 mM maltose and 1 mM 1,4-dithiothreitol, pH 8.0). The concentration of the protein was estimated by measuring absorbance at 280 nm (ε_280 nm_= 1,30,670 M^-1^cm^-1^) and samples were run on an SDS-PAGE gel to confirm the purity of the protein. The purified protein was further stored at 4 °C for future use.

The plasmid containing his_6_ tagged TEV protease was transformed into *E. coli* strain BL21(DE3) plysS. Cells were grown at 37 °C, 220 rpm, and overexpression was induced by 0.35 mM IPTG at 16 °C for 20 hours. Cultures were pelleted and stored at −80 °C for further use. The pellets were thawed at 30 °C and suspended in lysis buffer (25 mM HEPES, 150 mM NaCl, 20 mM KCl, and 20 mM MgCl_2_, pH 7.4) along with phenylmethylsulfonyl fluoride and lysozyme to enhance cell lysis which was carried out by probe sonication (5% amplitude, 15 sec ON/10 sec OFF for 25 minutes). The soluble fraction was separated by centrifugation and the supernatant was passed twice through a pre-equilibrated Ni-NTA column at 4 ^°^C. The beads were washed with wash buffer (lysis buffer + 20 mM imidazole) and protein was eluted with 300 mM imidazole and dialyzed against buffer without imidazole, overnight at 4 °C. Protein was concentrated using a 10 kDa MWCO filter and stored at −80 °C for further use.

### Fluorescence labeling

For labeling, purified FUS was concentrated using a 50 kDa MWCO amicon filter, and incubated with 0.3 mM tris(2-carboxyethyl)phosphine (TCEP) for 30 minutes on ice following which reactions with 1:4.5 molar ratio of protein: dye (for AlexaFluor488-C5-maleimide) and 1:30 (for Fluorescein-5-maleimide) were set up in native buffer at 25°C and kept under shaking for 2 hours in dark. Unreacted dye was removed by buffer exchange using 50 kDa MWCO amicon filters. Labeling efficiency was calculated by measuring absorbance at 280 nm (ε_280 nm_=1,30,670 M^-1^cm^-1^, for full-length FUS) and 494 nm (ε_494_ = 72,000 M^-1^cm^-1^, for AlexaFluor488-C5-maleimide and ε_494_ = 68,000 M^-1^cm^-1^, for F-5-M) to estimate total protein concentration and labeled protein concentration.

### Phase separation assays

Phase separation of FUS was initiated by TEV cleavage in a 1:10 molar ratio (TEV: protein) at room temperature in 20 mM sodium phosphate, pH 7.4. Turbidity of phase-separated samples was then estimated using 96-well NUNC optical bottom plates (Thermo Scientific) on a Multiskan Go (Thermo Scientific) plate reader by recording the absorbance at 350 nm. The protein concentration for all the experiments was fixed to 20 μM along with 0.1nM of iodide-modified silver nanoparticles Ag IMNPs (for reactions in the presence of nanoparticles). For phase separation in the presence of RNA, LLPS was induced in the presence of 25 ng/μL, 50 ng/μL, and 100 ng/μL polyU RNA with or without 0.1 nM Ag IMNPs. The total sample volume used was 100 μL for all the measurements and then background subtracted turbidity was plotted using Origin.

### Confocal microscopy

Confocal fluorescence imaging of FUS droplets with and without Ag IMNPs was performed on ZEISS LSM 980 Elyra 7 super-resolution microscope using a 63x oil-immersion objective (Numerical aperture 1.4). For visualizing droplets of FUS, 200 nM (1 %) of Alexa488 or F-5-M labeled protein was doped with unlabeled protein, and 2-3 μL of the freshly phase-separated sample was placed into a chamber made on a glass slide (Fisher Scientific 3” × 1” × 1 mm). The chamber made by using double-sided tape was then sealed with a square coverslip to avoid evaporation of the sample. For visualization of encapsulated Ag IMNPs (0.1nM), Alexa488-labeled protein was imaged using a 488-nm laser diode (11.9 mW), and Ag IMNPs were imaged using a 405-nm laser diode (11.9 mW). For images captured through the eyepiece, a metal halide lamp was used to excite both labeled protein and nanoparticles. All the confocal images were then processed and analyzed using ImageJ (NIH, Bethesda, USA).

### Fluorescence recovery after photobleaching (FRAP) measurements

FRAP experiments for droplets with and without Ag IMNPs were performed on ZEISS LSM 980 Elyra 7 super-resolution microscope using a 63x oil-immersion objective (Numerical aperture 1.4). All the FRAP experiments were performed using 200 nM (1%) of Alexa488-labeled protein. The recovery of the chosen region of interest (ROI) after photobleaching using a 488-nm laser was then recorded using ZEN Pro 2011 (ZEISS) software provided with the instrument. The fluorescence recovery curves were then normalized and plotted after background correction using Origin.

### Sedimentation assays

The absence of MBP within FUS droplets was confirmed using sedimentation assay. The MBP-FUS was cleaved by TEV protease to induce phase separation and after 20 minutes the reaction was centrifuged at 25,000 x g, 25 °C for 30 minutes to pellet down all the droplets (condensed phase). Both supernatant and the pellet were then separated carefully and the pellet was dissolved in 8 M urea. Samples were then heated and run on 12% SDS-PAGE along with the respective controls.

### Preparation of silver nanoparticles

Silver nanoparticles were prepared by the Lee-Meisel method as described previously^48^. Initially, 8.49 mg of silver nitrate was dissolved in 50 mL of filtered milli-Q water and stirred vigorously (1000 rpm) at its boiling point (∼ 98 ^°^C) for 30 minutes. One and five-tenths milliliters of freshly prepared 1% (w/v) aqueous trisodium citrate was added to the reaction mixture drop-wise and further stirred for additional 30 minutes till the color changed to yellow-green. The solution was cooled down to room temperature and was further characterized using UV-visible absorption spectroscopy, transmission electron microscopy, and zeta (*ζ*) potential measurements.

### Preparation of iodide-modified silver nanoparticles

One milliliter of silver nanoparticles was centrifuged in a 1.5 mL microcentrifuge tube at 5000 rpm for 15 minutes at room temperature. The supernatant was discarded and the resulting colloid was resuspended in 1 mL of Milli-Q water and centrifuged again. The resulting colloidal suspension (50 μL) was then mixed with an equal volume of 12 mM potassium iodide (KI) and incubated for 24 h at room temperature in dark. After incubation, the resulting iodide-modified nanoparticles were centrifuged at 5000 rpm for 10 minutes at room temperature and resuspended in 100 μL of milli-Q water. Resulting Ag IMNPs were characterized using UV-visible absorption spectroscopy, transmission electron microscopy, and zeta (*ζ*) potential measurements.

### UV-visible absorption spectroscopy

All the UV-vis absorption spectra were collected on a Multiskan Go (Thermo Scientific) plate reader using 96-well NUNC optical bottom plates (Thermo Scientific). The total sample volume used was 100 μL for all the measurements. Twenty micromolar full-length FUS was used whereas the concentration of Ag IMNPs used was fixed to 0.1 nM and 25 ng/μL, 50 ng/μL, and 100 ng/μL of poly-U RNA were used. Background subtracted absorption spectra from 300-800 nm were normalized and plotted using Origin.

### Dynamic light scattering (DLS)

Zeta potential measurements for iodide-modified nanoparticles were carried out on a Malvern Zetasizer Nano ZS90 instrument (Malvern, UK) using a He-Ne laser (632 nm) as an excitation source. All the measurements were carried out at room temperature and 0.05 nM of Ag IMNPs in filtered milli-Q water was used for estimating zeta potential.

### Transmission electron microscopy (TEM)

TEM images were obtained on a Jeol JEM-F200. Three microliters of half-diluted colloidal suspension were adsorbed on a 300-mesh carbon-coated electron microscopy grid and allowed to dry overnight. Histogram for nanoparticle size distribution was created using ImageJ (NIH, Bethesda, USA) software and plotted using Origin.

### Normal Raman and single-particle surface-enhanced Raman spectroscopy (SERS)

Single-droplet normal Raman and SERS spectra were recorded on an inVia laser Raman microscope (Renishaw, UK) at ∼ 25 ^°^C. For single-droplet normal Raman measurements droplet reaction (2 μL) of full-length FUS (20 μM) with or without RNA was placed on a glass slide covered with an aluminum foil and single droplets were focused using a 100x long working distance objective lens (Nikon, Japan). An NIR laser (785 nm) with an exposure time of 10 sec and 500 mW (100 %) laser power was used to excite the samples. Raman scattered light was collected and dispersed using a diffraction grating (1200 lines/mm) and was further detected by an air-cooled CCD detector whereas the Rayleigh scattered light was blocked using an edge filter of 785 nm. For single-droplet SERS measurements, phase separation of full-length FUS (20 μM) was set up in the presence of 0.1 nM Ag IMNPs with or without RNA, and 2 μL of the droplet reaction was placed on a glass slide covered with an aluminum foil. Single droplets were focused using a 50x long-working-distance objective lens (Nikon, Japan) and an NIR laser (785 nm) with an exposure time of 10 sec and 5 mW (1 %) laser power was used to excite the samples. Experiments were repeated with different batches of freshly purified protein and freshly prepared nanoparticles. Data was acquired using Wire 3.4 software provided with the Raman spectrometer. The collected Raman spectra were baseline corrected using cubic spline interpolation method and smoothened using Wire 3.4 software and plotted using Origin.

## Acknowledgments

We thank IISER Mohali, Department of Science and Technology (Nano-Mission to S.M. and FIST grant to the Department of Biological Sciences, IISER Mohali), Science and Engineering Research Board (SUPRA grant SPR/2020/000333 to S.M.), Ministry of Education, Govt. of India (Centre of Excellence grant to S.M.) for financial support, Prof. Dorothee Dormann (Ludwig-Maximilians University and Institute für Molekulare Biology, Mainz) for her kind gift of MBP-*Tev-*FUS-*Tev-*His_6_ plasmid, the TEM facility at IISER Mohali for electron microscopy, and Dr. Mily Bhattacharya (Thapar Institute) and the members of the Mukhopadhyay lab for critically reading this manuscript.

## Authors contributions

S.M. conceived the project. A.A., A.J., and S.M. further developed the concept and the experimental design. A.A., A.J. A.W., and S.G.P. performed the experiments and analyses. A.A. and A.J. prepared the figures and wrote the first draft. S.M. supervised the work, edited the manuscript, obtained funding, and provided the overall direction. All authors discussed the results and commented on the manuscript.

## Data availability

The data are available within the Article, Supplementary Information, and Source Data file.

