## Supplementary Information for "Single-Droplet Surface-Enhanced Raman Scattering Decodes the Molecular Language of Liquid-Liquid Phase Separation"

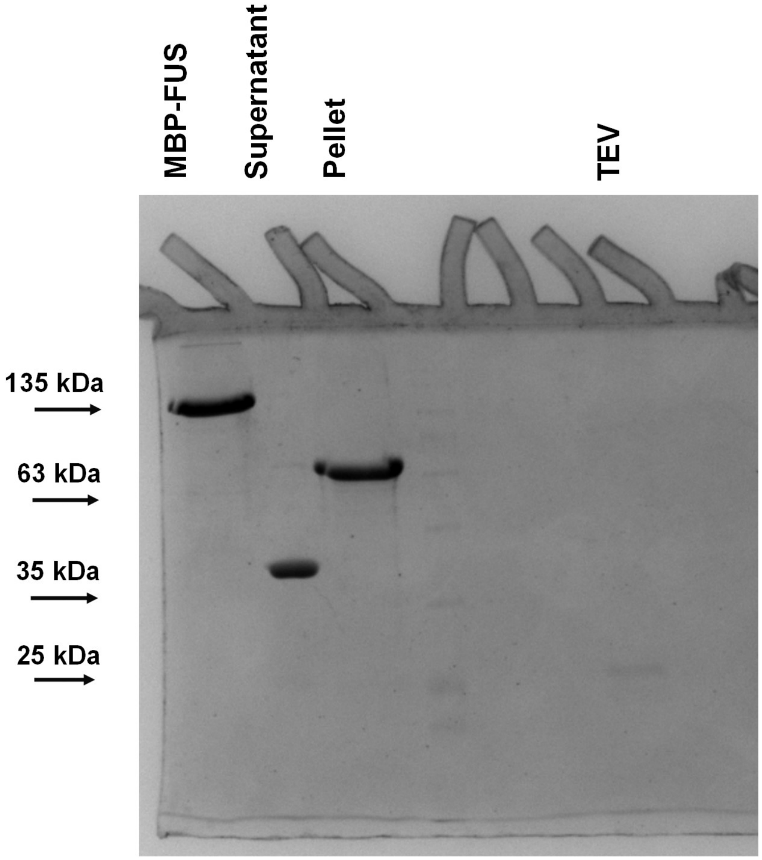


**Supplementary Figure 1:** SDS-PAGE (12%) depicting that the condensed phase is devoid of the maltose-binding protein (MBP) tag.

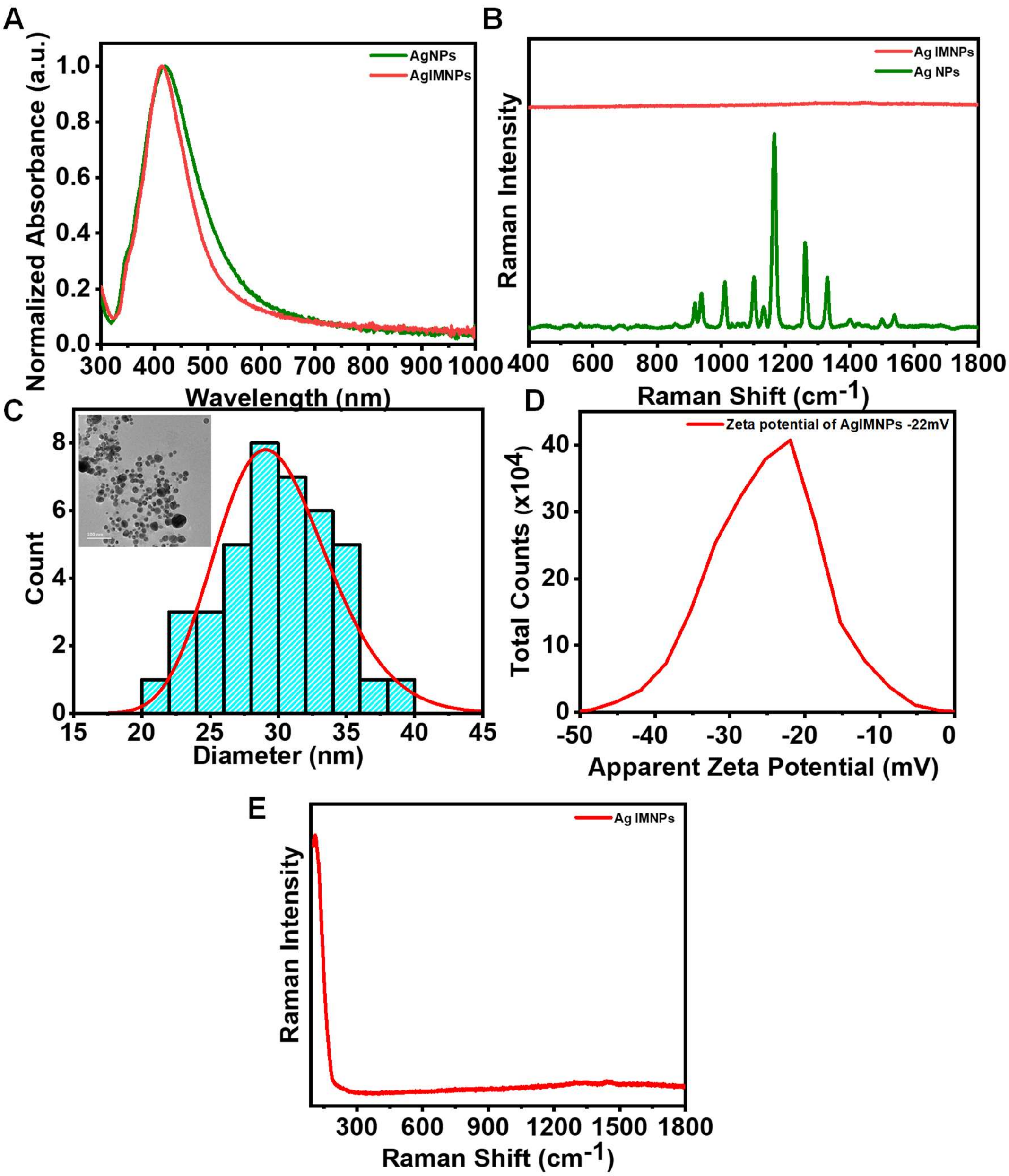


**Supplementary Figure 2: Preparation and characterization of iodide-modified silver nanoparticles (Ag IMNPs).** (A) UV-visible absorption spectra for silver nanoparticles (Ag NPs) (olive) and iodide-modified silver nanoparticles (Ag IMNPs) (red). (B) Raman spectra for silver nanoparticles (Ag NPs) (olive) and Ag IMNPs (red). (C) Histogram for nanoparticles size distribution derived from the TEM analysis. Size analysis performed by considering sizes of 40 different nanoparticles. Inset shows TEM image of Ag IMNPs. Scale bar: 100 nm. (D) Zeta-potential for Ag IMNPs. Plotted here is the mean of three different measurements. (E) SERS spectrum of Ag IMNPs corresponding to the Ag-I bond at 110 $cm^{-1}$.

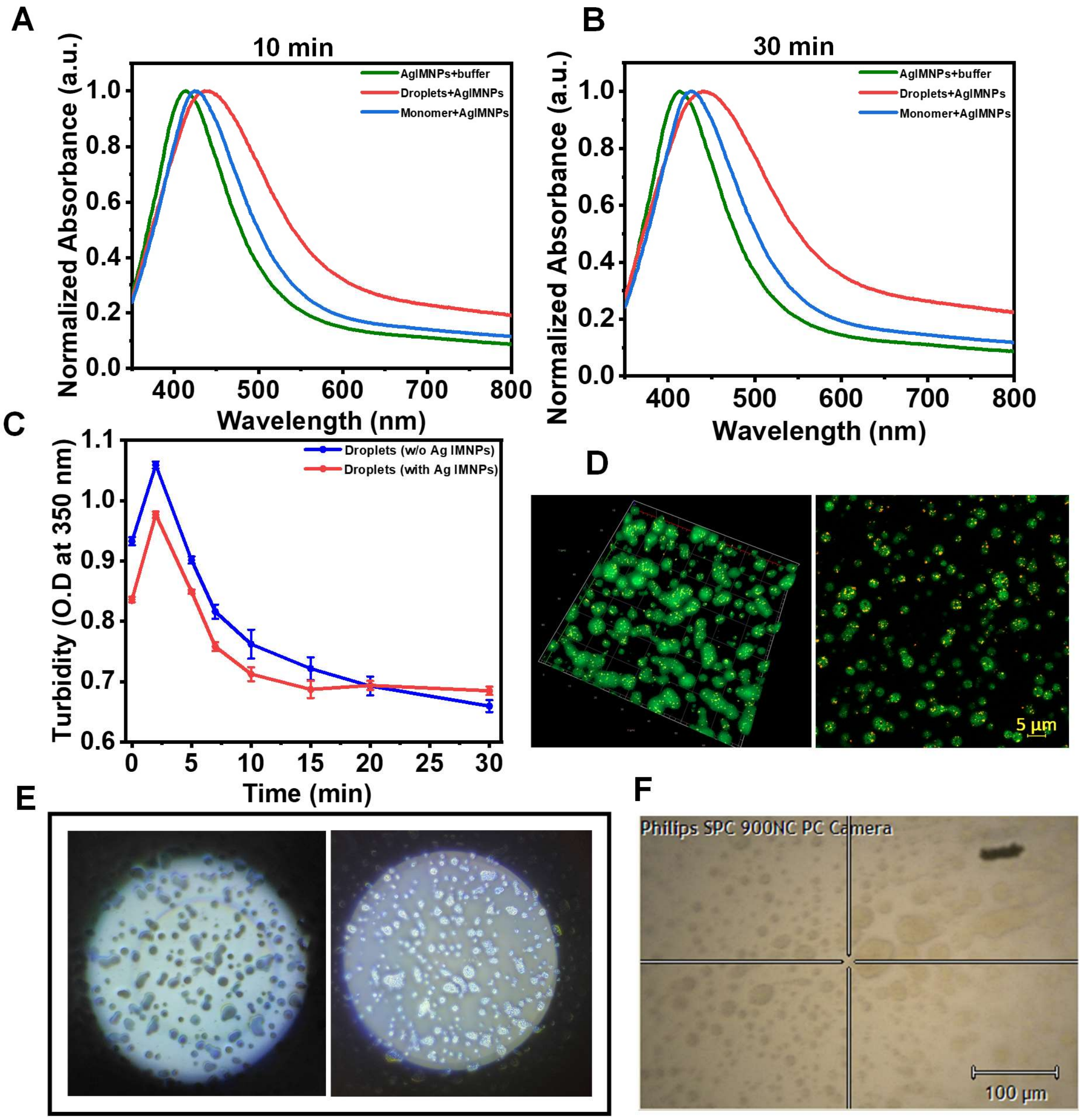


**Supplementary Figure 3: Interaction of FUS with iodide-modified silver nanoparticles (Ag IMNPs).** (A,B) UV-Visible absorption spectra of Ag IMNPs in phosphate buffer (olive), monomeric FUS in the presence of Ag IMNPs (blue), FUS droplets in the presence of Ag IMNPs (red) at 10 minutes and 30 minutes. (C) Turbidity plot of FUS droplets in the absence (blue) and presence (red) of Ag IMNPs (mean ± SEM; n=4). (D) 3-dimentional confocal image showing the presence of Ag IMNPs within FUS droplets. Confocal microscopy image of fluorescein-5-maleimide labeled FUS and Ag IMNPs indicating encapsulation of Ag IMNPs within the droplets. (E) Eye-piece image for FUS droplets without and with Ag IMNPs. (F) Encapsulation of Ag IMNPs as seen through a Raman microscope using 50x objective lens.

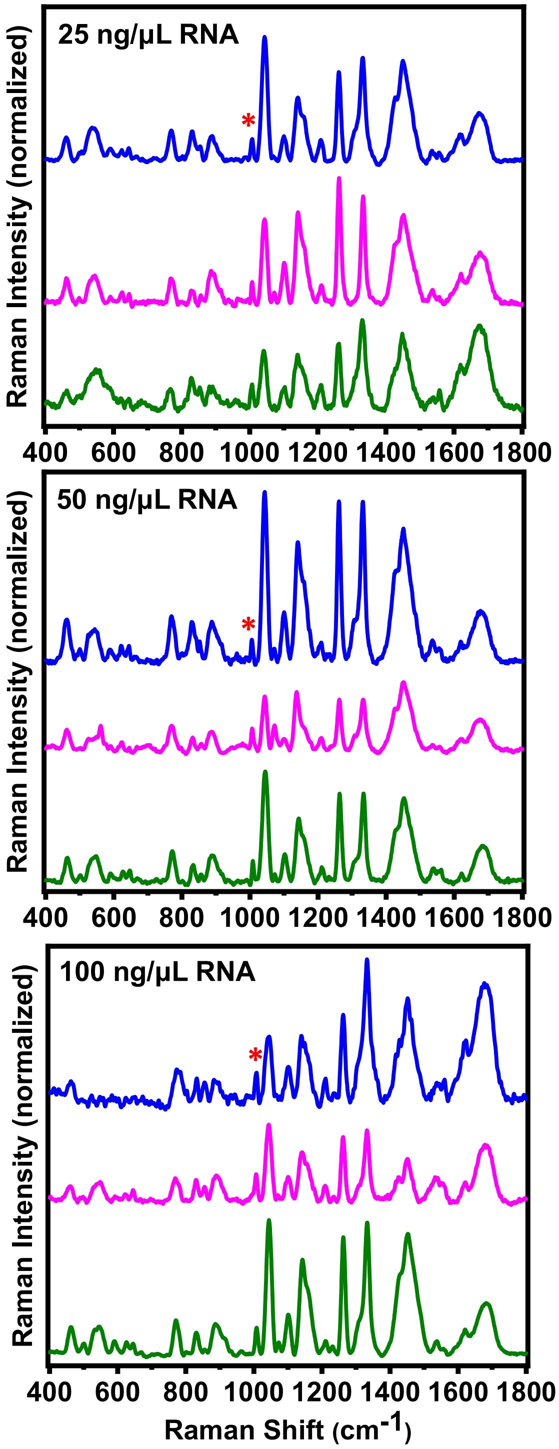


**Supplementary Figure 4:** Stacked single-droplet normal Raman spectra of FUS droplets at different concentrations of RNA (25 ng/µL, 50 ng/µL, and 100 ng/µL) (spectra recorded at 500 mW laser power with a 100x objective; 10 accumulations and 10 sec exposure time; number of droplets, n = 3).

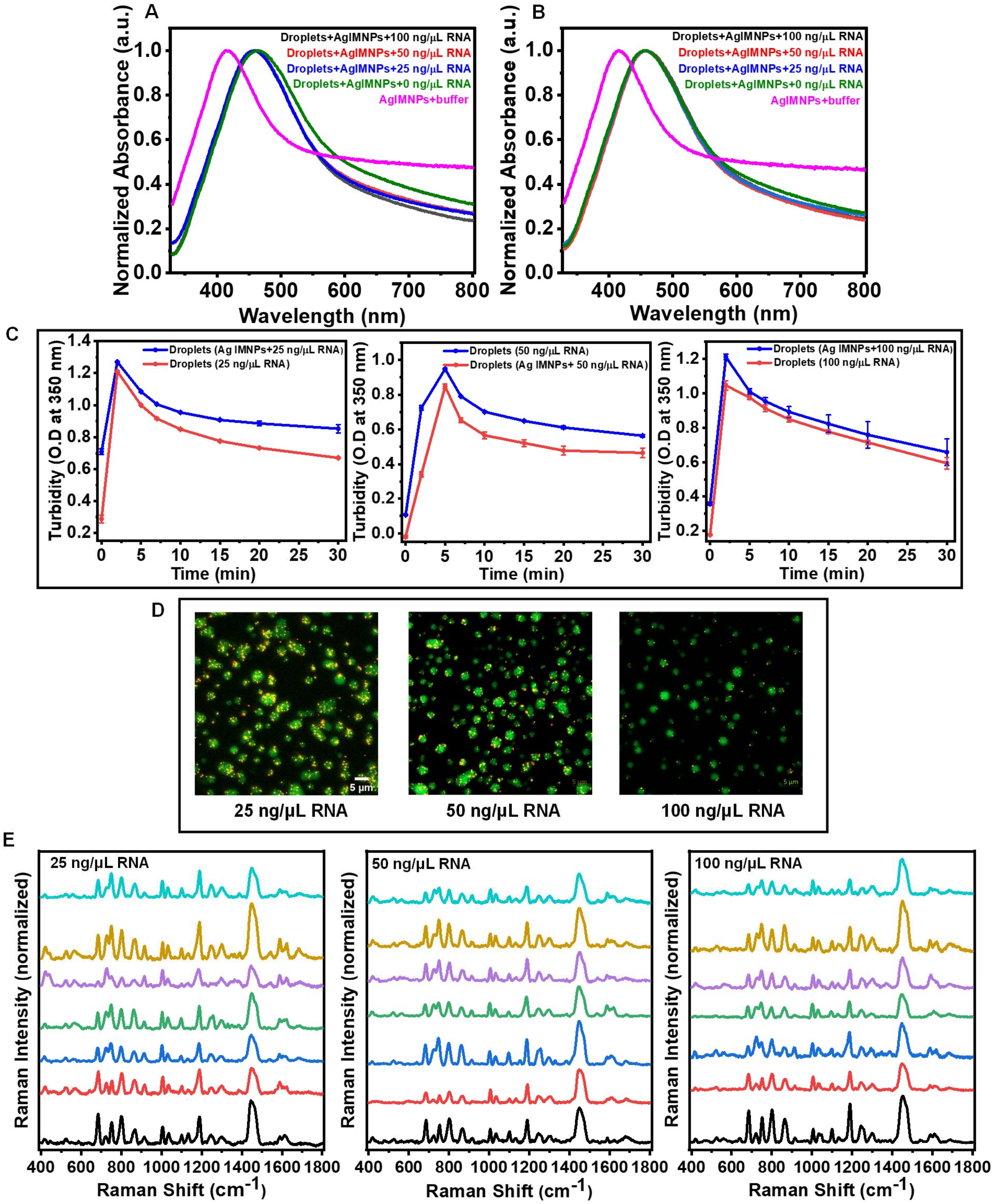


**Supplementary Figure 5: FUS droplets in the presence of Ag IMNPs and RNA.** (A,B) UV-Visible absorption spectra of Ag IMNPs in phosphate buffer (magenta) and FUS droplets at different concentrations of RNA (0 ng/ μL, 25 ng/μL, 50 ng/μL, and 100 ng/μL) in the presence of Ag IMNPs at 10 minutes and 30 minutes. (C) Solution turbidity plot of FUS droplets for different concentration of RNA (25 ng/μL, 50ng/μL, and 100 ng/μL) in the absence and presence of Ag IMNPs (mean ± SEM; n= 4). (D) Confocal microscopy images of FUS droplets for different concentrations of RNA (25 ng/µL, 50 ng/µL, and 100 ng/µL) in the presence of Ag IMNPs. (E) Stacked SERS spectra of FUS droplets at different concentrations of RNA (25 ng/µL, 50 ng/µL, and 100 ng/µL) (spectra recorded at 5 mW laser power with a 50x objective; 10 accumulations and 10 sec exposure time; number of droplets, n = 7).

**Supplementary Table 1.** Percentage analysis of δ(NH)-guanidinium moiety of arginine residues and $CH_2/CH_3$ deformation modes obtained after deconvolution of the region 1420-1490 $cm^{-1}$ for the single-droplet SERS spectra for various concentrations of RNA (0, 50, and 100 ng/µL).

| FUS droplets with Ag IMNPs with RNA | **δNH; Guanidinium moiety (1447 $cm^{-1}$) (in %)** | **$CH_2/CH_3$ deformations (1468 $cm^{-1}$) (in %)** |
|---|---|---|
| 0 ng/µL RNA | 71 ± 13 | 44 ± 13 |
| 50 ng/µL RNA | 68 ± 8 | 27 ± 8 |
| 100 ng/µL RNA | 60 ± 10 | 38 ± 10 |

**Supplementary Table 2.** Raman shift values and tentative band assignments[1-4] of normal Raman and SERS spectra of full-length FUS droplets.

| **Single-droplet Normal Raman** | **Single-droplet SERS** | **Peak assignments**[§] |
|---|---|---|
| 1673 (s) | - | Amide I (β-sheet) |
| 1621 (s) | 1621 (s) | Tyr (R stretch) |
| - | 1588 (s) | Phe, Trp, His |
| 1557 (m) | - | Trp |
| 1537 (w) | 1538 (w) | Trp, Amide II |
| - | 1447 (s) | δ(NH)-guanidinium moiety |
| 1451 (s) | - | δ($CH_2$/$CH_3$) |
| - | 1388 (w) | Asp, Glu $\nu_{sy}$[$COO^-$] |
| 1332 (s) | - | Trp, δ(CαH) |
| - | 1298 (s) | Amide III (α-helix) |
| 1262 (s) | - | Amide III (nonregular/turns) |
| - | 1246 (s) | Amide III (β-sheet) |
| 1209 (s) | 1213 (w) | Tyr [ν(C-C)] |
| - | 1188 (s) | Tyr, Phe, ν(C-N) |
| 1140 (s) | 1132 (m) | $\nu_{as}$(CαCN) |
| 1101 (s) | 1098 (s) | ν(C-C), ν(C-O), ν (C-N) |
| 1042 (s) | 1032 (s) | Phe [δ(R(CH))] |
| 1006 (s) | 1003 (s) | Phe R breathing |
| 958 (w) | - | ν(N-Cα-C) skeletal |
| - | 938 (s) | Backbone skeletal α-helix |
| - | 915 (s) | ν($COO^-$), C-C stretch of Pro ring |
| 890 (s) | - | Trp (N-H bend) |
| 857 (s) | 862 (s) | Tyr Fermi doublet |
| 830 (s) | - | Tyr Fermi doublet |
| 798 (w) | 800 (s) | ν(C-H), δ(N-H), Met [$\nu_{as}$(C-S-C)] |
| 767 (s) | 749 (s) | Trp [δ($R_{breathing}$)] |
| - | 724 (s) | Met [ν(C-S)] |
| - | 683 (s) | Met [ν(C-S)], δ(C-H) |

| | | |
|---|---|---|
| 646 (m) | 640 (w) | Tyr [γ(C-C)] |
| 623 (w) | - | Tyr, ν(C-S) |
| 597 (m) | - | δ($COO^-$) |
| - | 562 (m) | ν(S-S) |
| 539 (s) | 522 (m) | δ(skeletal), δ(N-H), ν(S-S) |
| 461 (s) | - | ν(C-S) |
| - | 419 (m) | Trp |

§δ, bending; ν, stretching; R, benzene ring; as, asymmetric; sy, symmetric; w, weak; m, medium; s, strong
